# Prediction of Plastic Degrading Microbes

**DOI:** 10.1101/2021.08.01.454681

**Authors:** N Hemalatha, Akhil Wilson, T Akhil

## Abstract

Plastic pollution is one of the challenging problems in the environment. But a life without plastic we cannot imagine. This paper deals with the prediction of plastic degrading microbes using Machine Learning. Here we have used Decision Tree, Random Forest, Support vector Machine and K Nearest Neighbor algorithms in order to predict the plastic degrading microbes. Among the four classifiers, Random Forest model gave the best accuracy of 99.1%.

## 1 Introduction

Plastic is nothing but a polymeric material. Plastic pollution has become one of the most stressing environmental issue, as rapidly increasing production of disposable plastic products overwhelms the world’s ability to deal with them. We cannot think of a life without plastic. Plastics revolutionized medicine with life-saving devices, made space travel possible, lightened cars, jets saved fuel and pollution, also saved lives with helmets, incubators, equipment’s for clean drinking water but on the other side it makes lot of environmental pollution. Practically it is difficult to avoid plastic completely from our daily life. The only solution to control the plastic pollution is degrading the plastic products rather than throwing it into the surroundings. Using the proper management, we can reduce the pollutions in the environment than what plastic creates.

*Streptococcus, Micrococcus, Staphylococcus, Moraxella, Psedomonas* these are the some of the plastic degrading microbes found in Indian mangrove soil. This outcome was a result of Japanese scientists in the year 2016 in which they found that a bacterium can easily break the plastic *polyethylene terephthalate* (PET). Further, they also found another bacterium called *Ideonella sakaiensia* obtained from the genus *Ideonella* and from the family *Comamonadaceae* which can break the plastic *polyethylene terephthalate* (PET). Once the bacterium acts, PET gets broken down into two i.e., ethylene glycol and DMT which can be used to create other materials. In this paper we have worked on developing a computational prediction tool with regard to plastic degrading protein sequences. Also have developed a database which consists of protein sequences collected from previous works which can degrade plastics. This research paper has been distributed like this: part 2 represents the different materials and also the methodology used to carry out this work. Then followed by the results and discussions in part 3 and Paper is concluded in part 4.

## 2 Methodology

This section describes the dataset used, different algorithms and features used for generating the computational tool.

### 2.1 Dataset

Plastic degrading protein sequences belonging to the alkB and CYP153 genes were cumulated from databases such as NCBI and UniprotKB. Around nine thousand positive protein sequences and six thousand negative sequences were obtained. For training data, we used the ratio 60:40 for positive and negative sequences.

### 2.2 Features Used

Six different features namely amino acid counts, dipeptide counts, amino acid ratio, hydrophobicity, hydrophilicity, acidity and basicity from microbial protein sequences were used for developing predicton. Features are explained in below sections.

#### 2.2.1 Amino acid count

In the amino acid count, we took the count of all the 20 amino acids in each microbial protein sequence making the dimension size 20 (Table 1). Equation used is

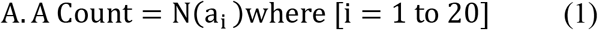

**Table 1.**
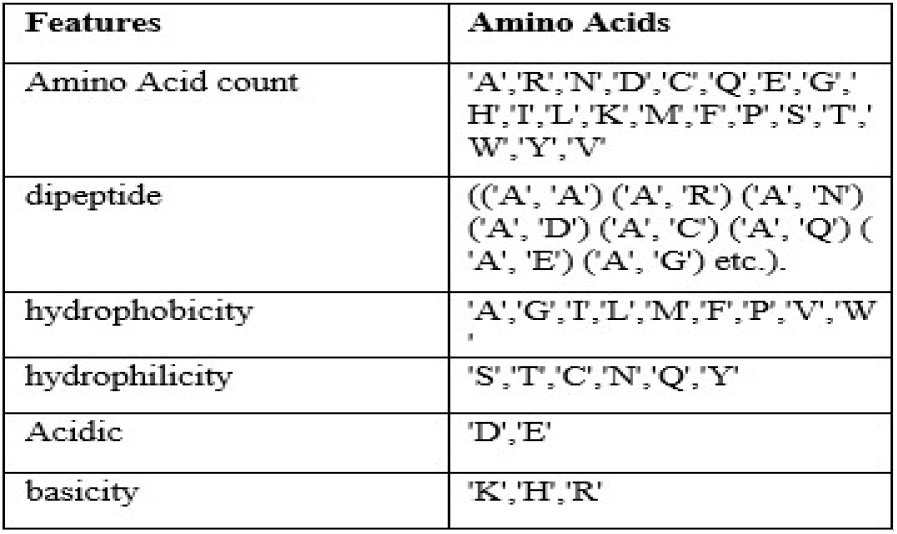
Details of the amino acids contained in different features

#### 2.2.2 Amino acid ratio

Here, count of each amino acid in a sequence was divided by the length of protein sequence. For this feature, feature dimension was 20 (Table 1). Equation used is

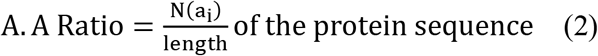

#### 2.2.3 Dipeptide count

In the dipeptide count, count of the occurrence of all dipeptide in the protein sequence was taken and dimension was of size 400 (Table 1).

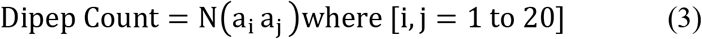

#### 2.2.4 Physiochemical properties

In this feature, hydrophobicity, hydrophilicity, acidity and basicity were considered (Table 1). Hydrophobicity had 9 amino acids and hence dimension of 6 whereas hydrophilicity six, acidity two and basicity had a dimension of three.

### 2.3 Data Pre-processing

Handling of the missing values is very important during the preprocessing as many machine learning algorithms do not support the missing data. The missing data can impact the performance of the model which is done through creating bias in the dataset. Hence, we can say that this bias can create a lack of relatability and trustworthiness in the dataset.

### 2.4 Algorithms used

In order to classify the plastic degrading microbes, we used the supervised machine learning techniques such as classification algorithms. Following subsections explains each classifier in detail.

#### 2.4.1 Decision trees

Decision Tree algorithm is a classification algorithm which has nodes and leaves. Nodes are split based on certain conditions and outcomes are obtained on the leaves. This is one of the most commonly using simplest classification algorithm.

#### 2.4.2 Random Forest

It is a classification algorithm in which it generates multiple decision trees based on the dataset. Each of the decision trees predicts the output and then finally it combines the outputs by voting.

#### 2.4.3 Support Vector Machine

This classifier creates a hyperplane on the dataset and then classifies the data points into classes.

#### 2.4.4 KNN (K-Nearest Neighbour)

It known as a lazy-Learner algorithm. This algorithms classifies the input values based on the similarity.

### 2.5 Performance measures

For measuring the performance of the computational prediction tool measures used were accuracy, confusion matrix, ROC-AUC and heat map which are described below.

#### 2.5.1 Accuracy

Accuracy can be defined as the ratio of number of the correct predictions to the total number of input values.

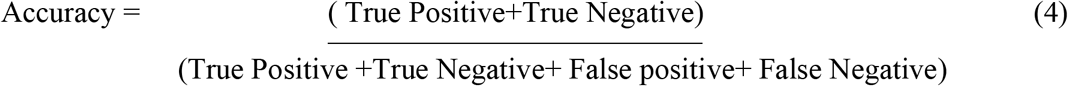

#### 2.5.2 Confusion matrix

Confusion Matrix which gives the output in the matrix form and explains the whole evaluation measures of the model which we created.

**Table 2.**
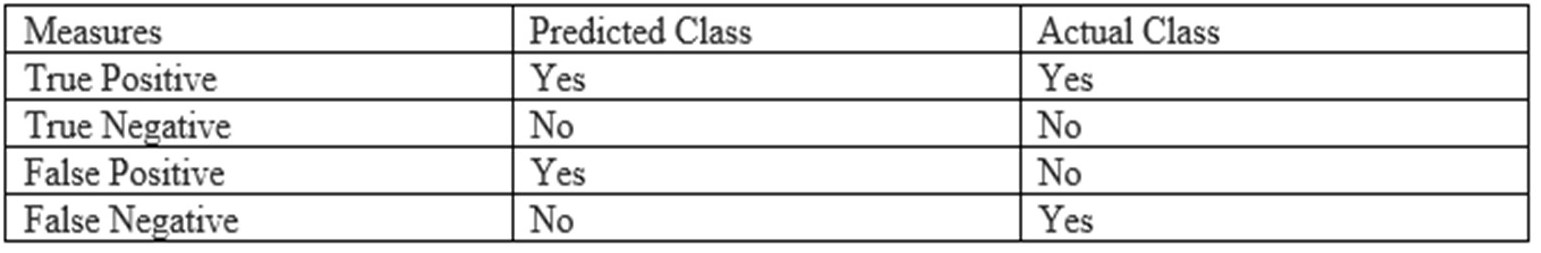
Measures of confusion matrix.

*F1 Score* we can define as a measurement of the models accuracy or harmonic mean of precision and the recall. This ranges from 0 to 1. It tells you that among the data how many instances it predicts correctly. The precision and recall can be calculated by using following equations.

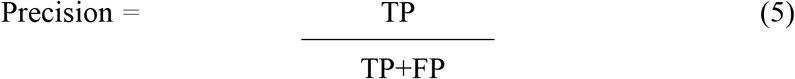

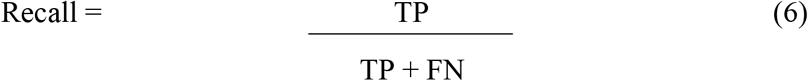

#### 2.5.3 ROC-AUC

AUC can be expanded as Area Under Curve. It is one of the metrics used for evaluation. As the value of the AUC rises the model performance for classification also rises. Before studying the AUC, we need to understand following measurements.

The True Positive Rate(TPR) we can defined as TP/ (FN+TP). It measures among all actual positive samples how many are correctly classified.

True Negative Rate(TNR) can be defined as TN / (FP+TN).

False Positive Rate(FPR) can be defined as FP / (FP+TN). It measures among all actual negative samples how many are incorrectly classified.

#### 2.5.4 Heat map

The heat map is nothing but a data visualization technique which helps to detect the correlation between the features.

### 2.6 Scikit learn

Scikit learn is a python library which gives many supervised and unsupervised learning algorithms. For this research work we have used NumPy, pandas and matplotlib from this library,

## 3. Results and Discussions

This section discusses the results obtained on pre-processing the dataset and then working the features of the dataset with four different machine learning classifiers.

### 3.1 Data pre-processing

In the present work, since twenty amino acids were compulsorily involved in all the plastic degrading microbes the chances of existence of the null values were nil. Using python environment, this was confirmed. Dataset used were also confirmed for skewness and existence of outliers. Details of all the three pre-processing are available in Table 3.

**Fig 1:**
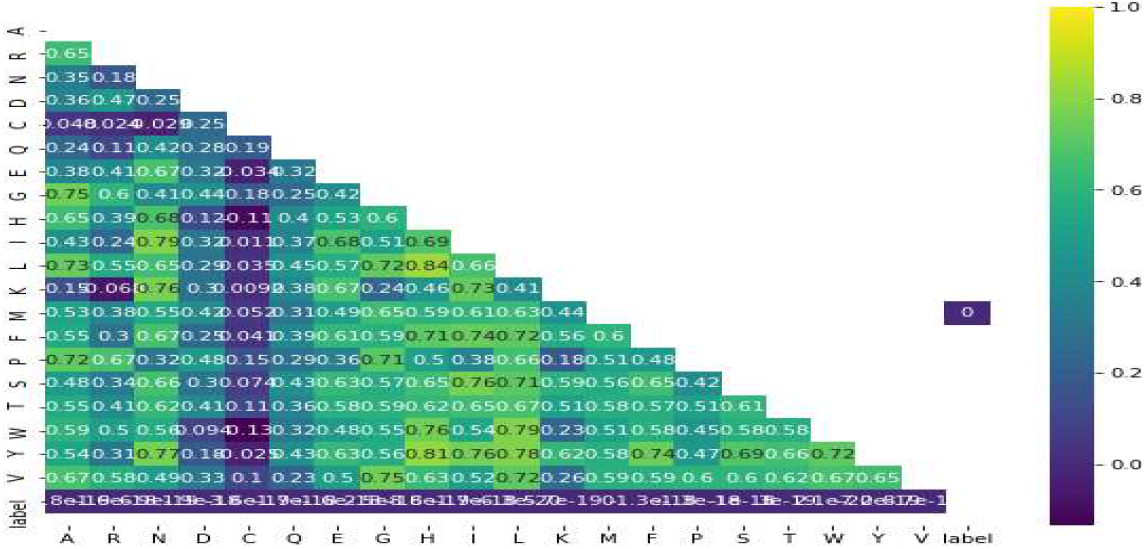
Heatmap

**Table 3:**
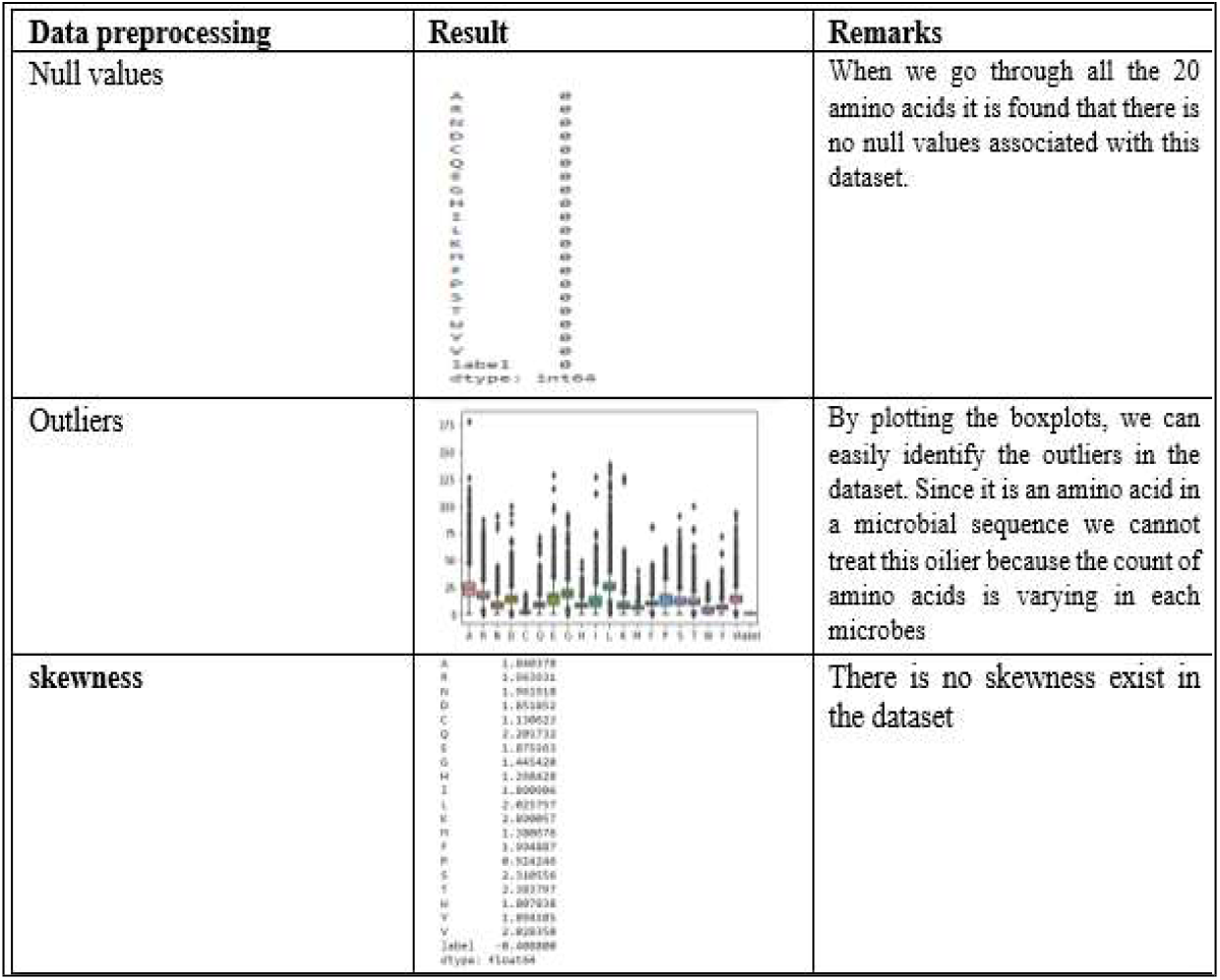
Table showing results of missing values, skewness and existence of outlier

### 3.2 Algorithm results

During experimentation of six features with four classifiers namely decision tree, Random Forest, SVM and KNN model obtained different accuracies which are listed in Table 4. From Table 5 it could be concluded that Random forest and KNN obtained best accuracy for the feature amino acid count. Diagrammatic representation is shown in Figure 2 and 3.

**Table 4:** Accuracies of different algorithms

| Model | Amino<br>Acid Co<br>unt | Dipeptid<br>e | hydroph<br>obic | Hydroph<br>ilic | Acidic | Basic | Amino<br>Acid Rat<br>io |
| --- | --- | --- | --- | --- | --- | --- | --- |
| RF | 99.1 | 12.088 | 98.433 | 93.733 | 89.2 | 84.866 | 98.766 |
| SVM | 98.7 | 36.55 | 96.133 | 88.7 | 89.0333 | 59.9 | 97.5 |
| DT | 98.33 | 12.08 | 97.166 | 90.5 | 89.16 | 83.7 | 98.3 |
| KNN | 99.1 | 26.88 | 98.266 | 90.733 | 89.0 | 83.666 | 99.2 |

**Table 5:** Accuracies in Amino acid count

| Model | accuracy |
| --- | --- |
| RF | 99.13333333333333 |
| SVM | 98.7 |
| DT | 98.33333333333333 |
| KNN | 99.1 |

**Fig 2:**
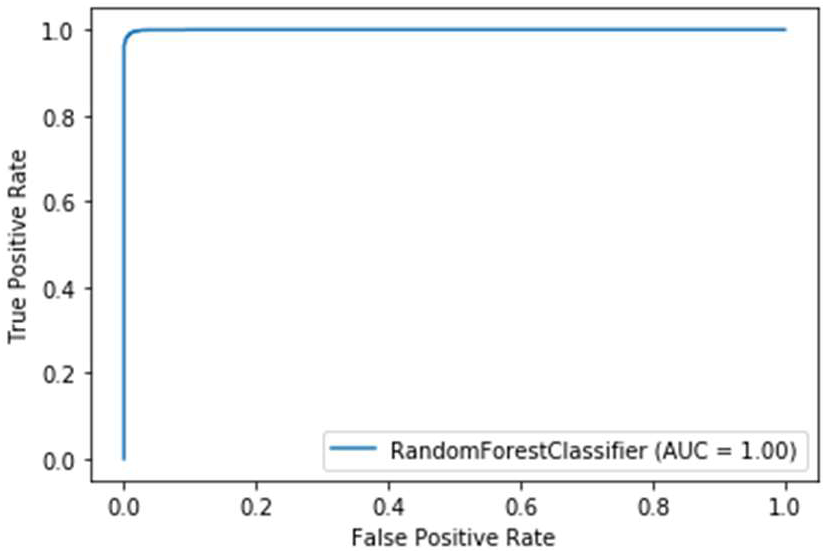
ROC-AUC curve of Random Forest for Amino acid count

**Fig 3:**
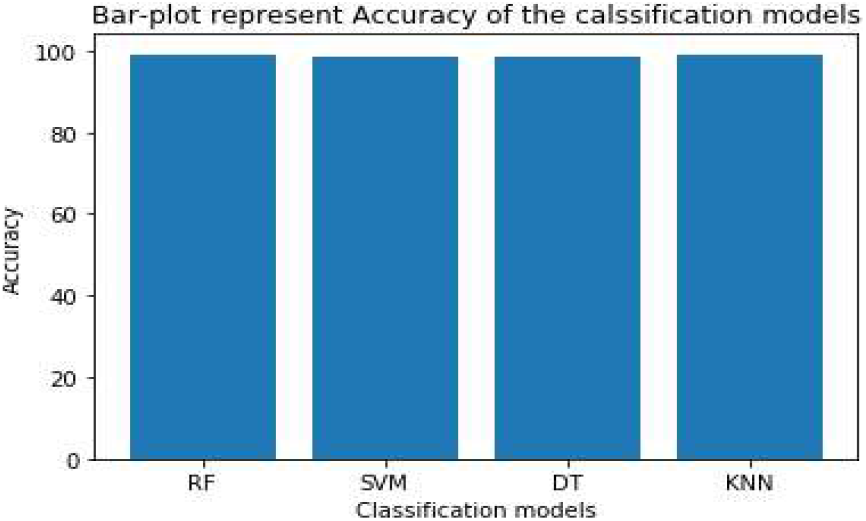
Bar diagram indicating the accuracies of 4 classifiers for amino acid count

## 4. Conclusion

In the present world to sustain without plastic is unimaginable but to overcome such situation only way out is to use microbes which can degrade the used plastic. In this work, we have attempted to develop a computational prediction tool which can classify a microbial protein if it is biodegradable or not. Four machine learning algorithms were used for this purpose along with six different features of p roteins. Out of the six features, it was found that the Amino acid count gave an accuracy of 99.1% with Random Forest and KNN classifiers. In the future work, we plan to develop a web server where we can host the prediction tool.

## Acknowledgement

This work is a part of the major research project funded by St. Aloysius Management.

## Notes

### Competing Interest Statement

All the three authors have no competing interest

